# Microbial Partner (MiPner) Analysis

**DOI:** 10.1101/2024.11.08.622289

**Authors:** Jeffrey L. Bennetzen, Josue Fernandez Canela, Vienna Elmgreen, Shaugnessy R. McCann, Mary E. Norris, Xiangyu Deng, Phillip Brailey-Jones

## Abstract

Although a few bacteria have been studied in great depth, relatively little is known about the characteristics of microbe-microbe interactions that occur within ecosystems on a daily basis. A simple, robust technique was developed to set up the foundation for investigating pairwise bacterial-bacterial interactions, using cell-cell binding as a self-selective mechanism to identify interesting bacterial species pairs. Using a Serratia marcescens strain (SMC43) isolated from Georgia soil as a “bait”, specific bacteria were purified by their specificity in binding SMC43 bacteria that were themselves attached to a wooden applicator stick. The isolated Microbial Partners (MiPners) were greatly enriched for members of the genera *Sphingobium* and Caulobacter. Two out of 24 streaked MiPners were unable to be separated from, and grow on the plate type tested without, SMC43. This suggests that the MiPner technology will be one strategy for purifying bacteria that were previously recalcitrant to culturing.

## 1 INTRODUCTION

All organisms pursue their life histories in the presence of other biological forms, some as competitors, some in a prey-predator relationship, some as co-operators, including symbionts. Microbes, in particular, commonly exist in complex communities, surrounded by hundreds to millions of other species of microbes, in such diverse and unstable environments as the atmosphere, bodies of water, multicellular hosts, and the soil. Despite this known ubiquitous and dynamic complexity, traditional microbiological research has concentrated on the study of one purified microbial species at a time to conduct hypothesis-based research, where the effects of a single variable are assayed in the presence of overall constancy. The advent and fabulous ongoing enrichment of “OMICS” technologies over the last 30 plus years (1–4) has provided a fully reversed perspective from the “one microbe at a time” strategy to an “everything at once” approach. This has facilitated a rapid expansion in the study of microbial ecology, especially to examine whole community microbiome interactions and to explore multi-taxa ecosystem-level interactions (5–7). The great wealth of data from OMICs approaches can provide deep and detailed correlations but come with their own weaknesses and limitations to interpretation. For instance, the enormous quantities of data and number of comparisons that are made always present statistical challenges, such as low resolving power and routine false positives, that must be resolved (8, 9). Hence, any correlations resulting from such analyses require further “hypothesis-driven” validation. With microbes, a more realistic environment for such confirmation experiments would require more than just the participation of a single microbial species.

In recent years, many research groups have attempted to create reproducible (that is, somewhat stable) microbial communities that are simpler versions of real-world assemblages (10–18). The problems that must be overcome include microbial competition, different growth rates, antimicrobials, incompatible metabolic properties, and dissimilar environmental requirements (19). We propose that a simpler type of relatively stable microbial community can be created with self-identifying microbial partners. The simplest of these partnerships, with two members, would be tremendously less complex than the real world, but much more complex than a single species experiment. In understanding how microbial species interact, it is unlikely that we will be able to fully conceptualize a complex natural environment until we begin to understand binary microbial interactions.

Such binary studies do exist, though many are consigned to the exploration of syntrophic interactions in co-culture (20). However, some have provided field-relevant and fascinating information regarding microbial competition as a tool for resistance against root diseases (21), for explaining how a soil bacterium can protect a soil fungus (22), or in the formation, function and/or destruction of biofilms (23–25). These previous examples all shared intense pursuits by dedicated research teams to investigate the biology of an “important” microbe. We believe that all microbes are worthy of investigation, and that the most important discoveries may come from microbes that we do not currently know anything about. Hence, a more general and facile method for identifying microbial partnerships would provide a useful step forward.

Here we present a novel method for isolating pairwise Microbial Partners (MiPners) from natural systems based on their propensity to physically associate with one another through microbe-microbe binding. Isolated partnerships can then be used to conduct hypothesis-based experiments. We have validated this method using a strain of *Serratia marcescens* isolated from Georgia soil, which was found to select and grow with only a tiny subset of the soil microbial collection, including with at least one bacterial strain that was unable to grow in the absence of its *S. marcescens* partner.

## 2 MATERIALS AND METHODS

### 2.1 Isolation and characterization of a *Serratia marcescens* strain, SMC43 (MiPner bait)

This study was designed to develop and demonstrate a technique for using a single microbial species as bait to identify and culture microbes that bind to- and can grow on plates with- a known bait microbe. Potential bait microbes were screened at the University of Georgia (UGA) as part of the course GENE4240L (“Experimental Microbiome Genetics”), which is designed to train students in the vagaries, certainties, and uncertainties of discovery science.

Through the Spring 2021 course, students isolated bacteria from soil samples collected on the UGA campus in Athens, GA (GPS coordinates: 33° 56’ 35.8’’ N, 83° 22’ 23.1’’ W). Soil suspensions were generated through mixing soil with phosphate buffered saline solution (PBS) from which 10^-2^, 10^-3^, and 10^-4^ dilutions were made with PBS. 100 microliters of each of the dilutions and the undiluted soil suspension was spread onto Soil Extract Agar (SEA, HiMedia Laboratories) plates. Plates were grown for two days at room temperature, and specific colonies of interest were isolated to pure culture through several rounds of streaking and re-culturing on Difco plates. Students Mary Norris and Molly Levin identified a red colony -later classified as *S. marcescens* strain C4-3 (hereafter referred to as SMC43)-which was chosen as the bait for a MiPner proof-of-concept study because of its red pigmentation (associated with production of the antimicrobial prodigiosin (26)) allowing for easily distinguishing the bait microbe from bound microbes.

DNA was extracted from an SMC43 overnight liquid culture using a protocol for high molecular weight DNA extraction (55). Libraries were prepared using the Rapid Barcoding kit (SQK-RBK004, Oxford Nanopore) and sequenced for 72 h using R9.4.1 flow cells (FLO-MIN106, Oxford Nanopore) on a GridION instrument. The genome was assembled with Canu (56). The contigs were circularized using Circlator (57) The assembly was taxonomically analyzed with TYGS (58). The genome was annotated using RAST (59). A BRIG chart was constructed and annotated using Proksee (60).

### 2.2 Initial conceptualization and implementation of the MiPner identification strategy

To isolate MiPners of SMC43, we conceived a series of steps that required both cell-cell binding of environmental bacteria to SMC43 and subsequent growth with SMC43 on Difco plates. The first component is a cultured microbial bait, in this case SMC43. The second component is a soil suspension from which MiPners can be captured through their physical associations with the bait microbe. A set of control experiments were also designed to ensure that cultured bacteria were indeed due to associations between field-sourced bacterial communities with the SMC43 bait, and not due to other factors, including lab contamination or natural binding capacity to the bait sticks used. The full MiPner and control experiment strategy is detailed in Fig. S1.

To prepare the SMC43 bait, overnight cultures were made from pure SMC43 colonies using Difco nutrient broth held at room temperature with 150 RPM shaking. Soil solutions were made by mixing soil collected in January 2023 from the UGA campus (GPS coordinates: 33° 55’ 44.9’’ N, 83° 21’ 46.1’’ W) with 0.1X Saline-Sodium Citrate solution (15 mM NaCl, 1.5 mM Sodium Citrate, pH 7.0) at room temperature. The mixture was shaken at 250 rpm for 10 minutes and left for a further 10 minutes for most particulate matter to settle at the bottom of the container. The upper 25% of this suspension was decanted into a sterile 500 ml container. This soil suspension served as the microbial inoculum source for both the MiPner and control experiments (Fig. S1A)

### 2.3 Control experiments for MiPner identification

To characterize and quantify the ‘source’ communities from which MiPners would be picked, 2 ml aliquots of soil suspension were repeatedly transferred to tubes and centrifuged at 10,000 g for 10 minutes to precipitate bacteria and other microbes. Once 250 mg of precipitate had been collected, DNA was extracted from the pellet using a DNAeasy Power Soil Kit following the manufacturers protocols (Qiagen, Hilden, Germany) (Fig S1.B). This collection process was carried out in triplicate to generate three technical replicate samples from the soil suspension. This is an important step in MiPner determination because bacteria found in the MiPner experiments but not in the source community may be from environmental contamination.

A second set of control experiments were carried out to identify how culturing methods and conditions could intrinsically bias the observed community when culturing both with and without SMC43. These experiments again act as controls to ensure that putative MiPners identified were indeed present within and culturable from the source community and not contaminants. For the first culture control (labeled ‘Pure’), the soil suspension was used as an inoculant for direct culturing on 0.1 X Difco plates. 50 ul of the suspension was pipetted onto each plate across three technical replicates and spread using sterile glass beads. Plates were left for three days at room temperature (approximately 22 °C). To capture the communities from the plates for DNA extraction, 1 ml of 0.1 X SSC was added to the plate and gently shaken for five seconds, poured off into a tube, and then used as input for DNA extraction (57) (Fig. S1C). All subsequent plates described were cultured and extracted in this manner unless otherwise specified.

The second culture control (Mixners) was used to identify which bacteria could survive or thrive in growth with SMC43 on 0.1X Difco plates. Two experiments were performed using different ratios of SMC43 and the soil solution inputs. For the MixnerA experiments, 500 μl of the soil suspension were mixed with 500 μl of an SMC43 overnight culture in a sterile 2 ml tube. 50 μl of this mixture was spread on each of three 0.1x Difco plates with glass beads. For the MixnerB experiments, 100 μl of the soil suspension was mixed with 900 μl of a SMC43 overnight culture. As before, 50 μl of the mixture was spread onto each of three 0.1x Difco plates for culturing and DNA extraction (Fig. S1E).

A final set of control experiments was conducted to identify microbes capable of directly binding to the applicator stick in the absence of SMC43 (referred to as ‘Binder’ experiments). Any bacteria that are able to do this would be unreliable MiPners if observed in such experiments, as we would be unable to distinguish whether they were observed due to their association with SMC43 or due to their natural binding capacity to the applicator stick. Sterile wooden applicators were set in a tube containing 1 ml of soil solution for 5 minutes, and afterwards were rinsed with 0.1X SSC and then dabbed onto a fresh Kimwipe to remove the excess liquid. Each of these applicators were used to streak onto 0.1X Difco plates for culturing and DNA extraction.

### 2.4 MiPner identification experiments

Following all control experiments, the MiPner experiment entailed immersing the sterile wooden applicators first in a tube containing one ml of an overnight SMC43 culture (in 0.1X Difco) for 5 minutes. After 5 minutes, the applicators were rinsed with 0.1X SSC and then dabbed onto a fresh Kimwipe to remove excess liquid. The applicators were then immersed in a tube with one ml of soil solution. After 5 minutes, each applicator was individually rinsed with 0.1X SSC and then dabbed onto a fresh Kimwipe to remove excess liquid. Each of these applicators was used to streak onto 0.1X Difco plates for culturing and DNA extraction. A demonstrative example of the MiPner experiment and ‘binder’ controls is given in Fig. S2.

### 2.5 Sequencing and data analysis of MiPner and control experiments

For sequencing library preparation, DNA concentration was measured fluorometrically and diluted to 2.5 ng/ul. Libraries were enzymatically fragmented and barcoded using the Plexwell 384 plate-based library preparation system (SeqWell, California), following the manufacturer’s protocol.

Prepared libraries were sequenced on a NovaSeqX flow cell lane in PE150 mode (Illumina, San Diego, CA USA). Raw sequencing reads were trimmed, and quality filtered using fastp v0.23 (61). Reads were classified using Kraken2 v2.1.3 (27) with a paired-end setup. Reads were classified against the standard Kraken2 database which draws genomes from the NCBI RefSeq database (downloaded: March 27, 2023). The relative abundances of sequences at both the genus and species level were then re-estimated using Bracken 2.7 (65). Bracken reports were merged using kraken-tools *v 1.2* (28), and archaeal, viral and human reads were removed from the bracken reports prior to summarizing genera and species abundances.

### 2.6 Sequencing and genome analysis of isolated colonies from MiPner experiments

DNA was extracted from isolated colonies with a protocol for high molecular weight DNA (55). Libraries were prepared using the Rapid Barcoding kit (SQK-RBK004, Oxford Nanopore) and sequenced for 72 h using R9.4.1 flow cells (FLO-MIN106, Oxford Nanopore) on a GridION instrument. The genomes were assembled with Canu (56). The assemblies were taxonomically analyzed with TYGS (58)

## 3 RESULTS

### 3.1 Isolation and characterization of a novel strain of S. marcescens to use as bait

Long-read sequence information indicated that the isolated red colony belongs to a strain of *S. marcescens*, named C4-3. This yielded an assembled chromosome of 5,092,593 bp with a 3,071 bp plasmid. Further annotation by the GENE4240L students who isolated this bacterium, Mary Norris, and Molly Levine, indicated that it carried a predicted 4939 protein-encoding sequences in the chromosome and one in the plasmid (Fig. S1).

### 3.2 Observed communities across the control experiments

The bacterial community observed in the soil sample was highly diverse, containing a mean genera richness of 681.7 ± 2.8 SE across the triplicate technical replicates. The dominant genera found within the soil were *Bradyrhizobium, Priestia* and *Strepromyces*, with a high abundance of low-frequency microbial groups (labelled as ‘Other’, Fig. 2A). On the 0.1X Difco plates, there were far fewer observed genera from culture (147.7 ± 4.5SE), of which very few dominated the whole community (i.e., the 10 highest abundant genera combined constituted ∼ 97% of all observed reads in the three replicates). The 0.1X Difco plates greatly enhanced growth of bacteria belonging to the genera *Enterobacter*, *Flavobacteria* and *Pseudomonas* at the expense of *Streptomyces*, *Priestia*, *Bradyrhizobium* and many other genera (Fig. 2B).

Culturing the soil bacteria in the presence of SMC43 was expected to act as an additional selective force on the culturable bacteria. Figure 3 provides the results of mixing the soil solution with two different ratios of SMC43, and then growing this mixture for 3 days on 0.1X Difco plates. These results indicate that growth with SMC43 affected microbial diversity dramatically. Whereas growth on the plate without excessive SMC43 competition yielded an abundance of *Enterobacter* (Fig. 2B), the SMC43-spiked plates enhanced the abundance of *Flavobacter*, *Janthinobacterium* and *Pseudomonas* (Fig. 3). Most Enterobacterial species were apparently killed or otherwise growth-inhibited by SMC43. The dosage of SMC43 also made a difference, with pedobacter and xanthomonads doing better at a lower SMC43 dosage. We use the generic name “Mixner” to denote these microbes that survive growth with our bait microbe, SMC43. Thus, Mixners could be partners in interactions with SMC43 even if they do not bind under our study conditions.

Some of the apparent MiPners might be merely microbes that stuck directly to the applicator stick, and not to SMC43. Our stick-binding results indicated that members of the genera *Enterobacter*, *Chryseobacterium* and *Pseudomonas* were particularly avid binders to the applicator stick under these conditions from our studied soil solution (Fig. 4). Through further observation of abundances at the species level (Supplementary Table S2), we can reliably state that numerous species of both *Enterobacter* and *Chryseobacterium* were highly prevalent stick-binders. While most *Psuedomonads* can bind to the stick to varying degrees of success, there are four taxa that display marked increases in their relative abundance when grown with SMC43.

### 3.3 MiPner identification and isolation

The observed community of MiPners varies across the triplicated technical replicates (Fig. 5), a presumed outcome of the samples that were streaked on the Difco plates containing very few bacteria that passed all the requirements, especially binding to SMC43 (see Fig 1, lines 3 and 4). In this case, stochastic effects could play an influential role in inter-replicate variability because of low depth in the sampling. Regardless, at least two samples contained major representation of the genera *Pseudomonas*, *Sphingobium*, and *Caulobacter* (Fig. 5). Despite its presence as the most abundant genus in the Mixner experiments (Fig. 3), *Flavobacterium*, was not among the ten highest abundant genera in these MiPner experiments, indicating a general inability to bind SMC43 under our study conditions.

**Figure 1.**
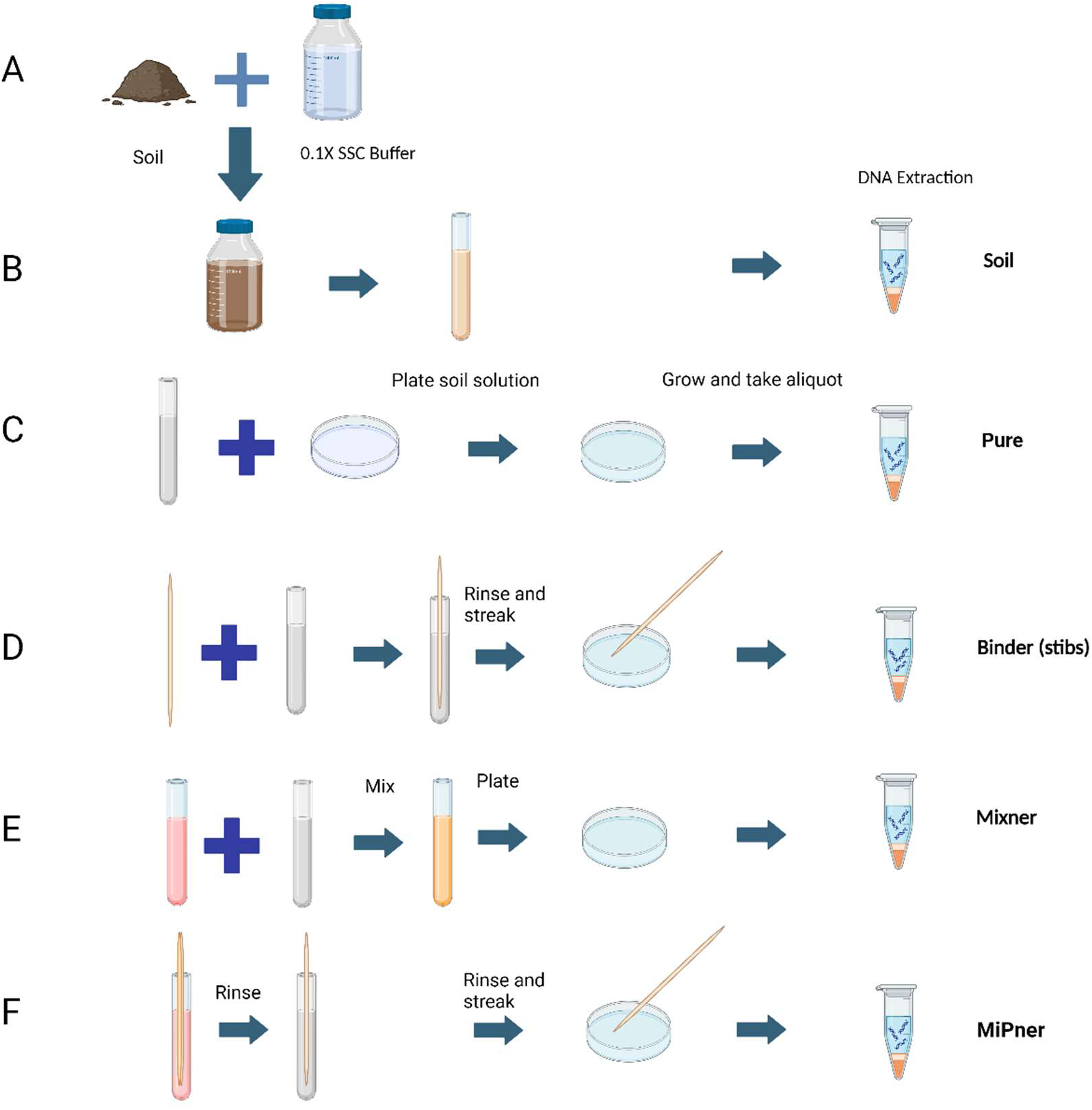
MiPner steps and controls. A) Starting soil and buffer; B) The soil suspension is created from mixing the soil and buffer, allowing this to settle, and aliquoting only the supernatant (Soil); C) Microbes are plate cultured from the soil suspension (Pure); D) A wooden applicator is submerged into the soil suspension and used for plate culturing of microbes (Binder); E) Microbes are plate cultured from the soil suspension mixed with SMC43 (Mixner); F) A wooden applicator is submerged into the SMC43 solution, followed by submersion in the soil suspension, and used for plate culturing of SMC43-bound microbes (MiPner).

### 3.4 Genus-level enrichments across MiPner and control experiments

At the genus level, no *Enterobacter*, *Pseudomonas* or *Cupriavidus* could be trusted as a true MiPner isolate because of their strong affinity (see the “Binder Average” column, Table 1) for binding to the applicator sticks that we used, despite being found in high abundance in the MiPner samples. In contrast, *Sphingobium* and *Caulobacter* were not seen to bind to the applicator, and were rare in all steps, indicating at least 100-fold relative enrichment compared to the starting soil suspension.

**Table 1.** Mean relative abundances per experimental condition of the ten highest abundances putative MiPner genera identified from the MiPner experiments. Relative abundances are displayed across both the MiPner experiments and associated control experiments.

| Ten Most Abundant MiPner Genera | MiPner Average (%) | Binder Average (%) | Mixner A Average (%) | Mixner B Average (%) | Soil Solution on Plate Average (%) | Soil Solution Average (%) |
| --- | --- | --- | --- | --- | --- | --- |
| *Pseudomonas* | 43.13 | 4.86 | 8.66 | 7.81 | 7.06 | 0.78 |
| *Sphingobium* | 27.19 | Not present | 1.08E-03 | 0.01 | 0.01 | 0.26 |
| *Caulobacter* | 15.76 | Not present | 4.24E-03 | 0.01 | 0.01 | 0.12 |
| *Massilia* | 3.96 | 0.16 | 5.99 | 3.76 | 0.32 | 0.28 |
| *Cupriavidus* | 3.17 | 0.91 | 0.29 | 1.13 | 1.99 | 3.65 |
| *Enterobacter* | 2.44 | 81.27 | 0.88 | 0.94 | 72.56 | 2.75 |
| *Duganella* | 0.96 | 0.10 | 1.13 | 3.32 | 0.49 | 0.06 |
| *Burkholderia* | 0.49 | 2.27E-03 | 0.03 | 0.08 | 0.04 | 0.57 |
| *Bradyrhizobium* | 0.38 | 3.76E-04 | 0.01 | 0.01 | 6.26E-04 | 8.17 |
| *Janthinobacterium* | 0.35 | 0.17 | 15.11 | 9.60 | 1.09 | 0.04 |

### 3.4 Species-level enrichments across MiPner and control experiments

At the species assignment-level, 33 individual taxa were identified belonging to the *Serratia* genus. *S. marcescens* was identified in expectedly high relative abundances across the experiments where SMC43 was added (MiPner: 64.24 % ± 6.35; MixNer E+F: 79.18 % ± 3.63, Supplementary Table S1), and in low abundances within the binder (0.01 % ± 0.01) and pure (0.09 % ± 0.004) experiments. *S. marcescens* was not observed to be present in any soil samples. The presence of *S. marcescens* in binder and pure samples could therefore be due to either incredibly low-level cross-contamination of these samples during the experiments and / or sequence prep, or likely due to levels of *S. marcescens* (and other related species) in the soil extracts that was below the detection threshold of sequencing in these samples but not within other experiments where diversity was artificially lowered by experimental conditions. Due to the nature of *Kraken2 / Bracken* taxonomy assignment and abundance estimation, the majority of the remaining *Serratia* taxa sequence assignments are also likely derived from low-information DNA regions of the sequenced SMC43 isolate or related environmental taxa. Of the 32 *Serratia* taxa not directly attributed to *S. marcescens,* only four were found in experiments where SMC43 was not added, and when they were recovered in these samples, they were in incredibly low abundance (< 0.01% mean relative abundance in any experiment group). This demonstrates that for mixed bacterial communities, the interpretation of species-level assignments based on *Kraken2 / Bracken* requires some caution.

All Serratia-related taxa were removed for subsequent analyses, so relative abundances of the remaining taxa were calculated without Serratia. The top 100 most abundant taxa across the MiPner experiments were considered to determine which individual taxa could be regarded as putative MiPners based on their presence in both the MiPner experiment and ‘Binder’ control experiments (Supplementary Table S2). 18 taxa were identified with > 1% relative abundance in at least one MiPner experiment sample, of which 6 were identified as putative MiPners, and 12 could not be assigned MiPner status due to their presence in the binder control experiments. Putative MiPners include *Spingobium yanoikuyae, Caulobacter segnis, Sphingobium sp. PAMC28499, Sphingobium sp. LF-16, Caulobacter vibrioides* and *Burkholderia thailandensis.* Of the 12 taxa which could not be confidently assigned MiPner status with high abundances, there were 6 *Pseudomonas,* 3 *Masilia, 1 Cupravidus, 1 Duganella*, and 1 *Enterobacter* assigned taxa.

21 additional putative MiPners were identified at a low abundance (< 1% relative abundance in at least one MiPner experiment sample). These included several taxa that were not considered when examining genus level associations including *Bordtella parapertussis, Diaphorobacter polyhydroxybutyrativorans, Nordella sp. HKS 07, Pectobacterium carotovorum, Lautropia mirabilis* and *Yersinia Pestis.* Additionally there were 5 *Caulobacter*, 4 *Sphingobium*, 2 *Bradyrhizobium,* 2 *Masilia* and 2 *Pseudomonas* taxa identified as putative MiPners. The detection of Pseudomonads as putative MiPners in contrast to our genus-level assessment shows that there may be some utility to considering a species level definition. As with all low-abundance taxa however, these two identified *Pseudomonas* MiPners are unlikely to be readily detectable through onward culturing, and when including the high abundance (> 1%) Pseudomonads detected represent only two out of 36 observed taxa in this genus. Low capacity for species resolution as demonstrated by our *Serratia* analysis also calls into question how definitive this approach can be in comparison to genus level assessments without onward culturing.

### 3.5 MiPner genome sequences and phylogenetic assessment

Our results demonstrate that the MiPner analysis excluded most microbes from the soil solution while selecting a tiny subset that could both bind SMC43 and grow with it under the conditions investigated. As can be seen (Fig. 4), *Pseudomonas*, *Sphingobium*, and *Caulobacter* were the most abundant genera represented in this MiPner experiment.

**Figure 2.**
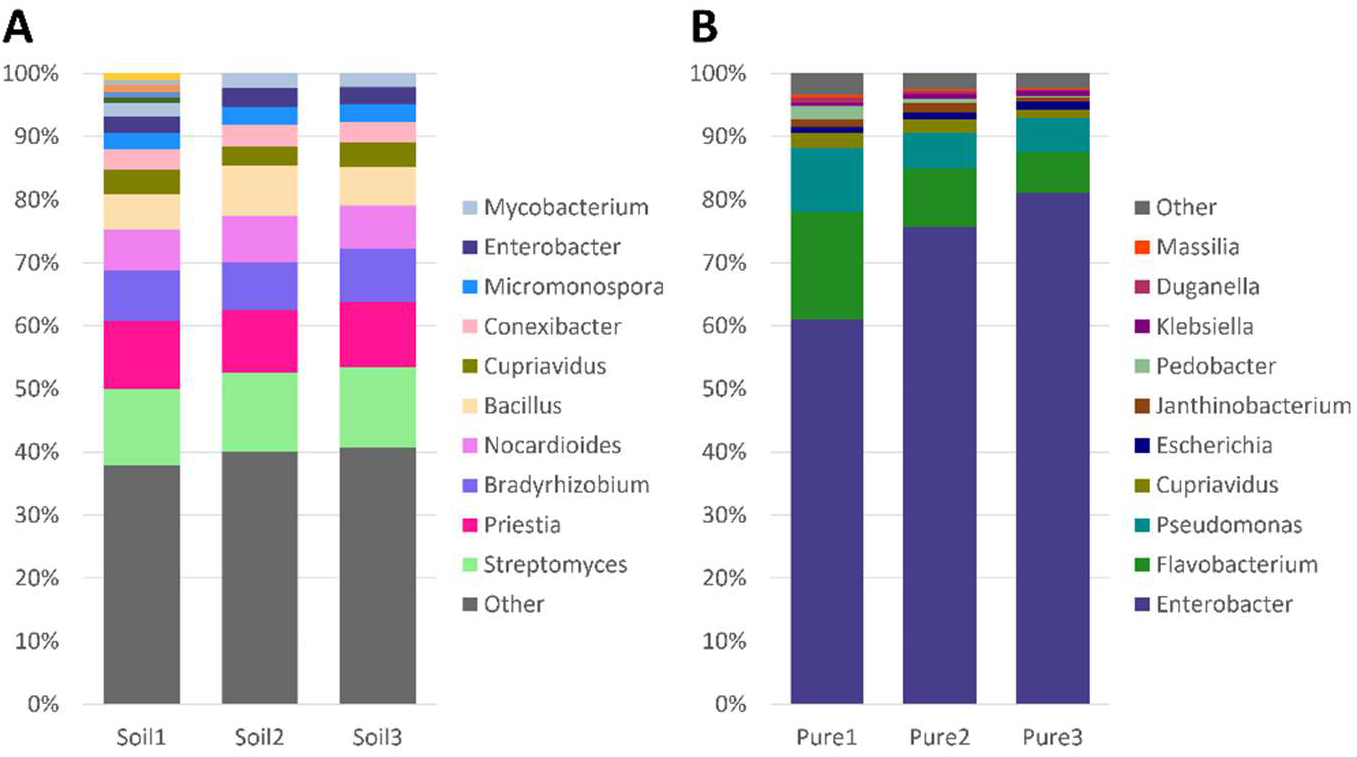
Taxa charts displaying the ten highest abundance bacterial genera of each treatment across from (A) the initial soil suspension and (B) the same suspension grown on 0.1X Difco plates for three days at room temperature. Genera are depicted in abundance order. Triplicate technical replicates are shown for each treatment.

**Figure 3.**
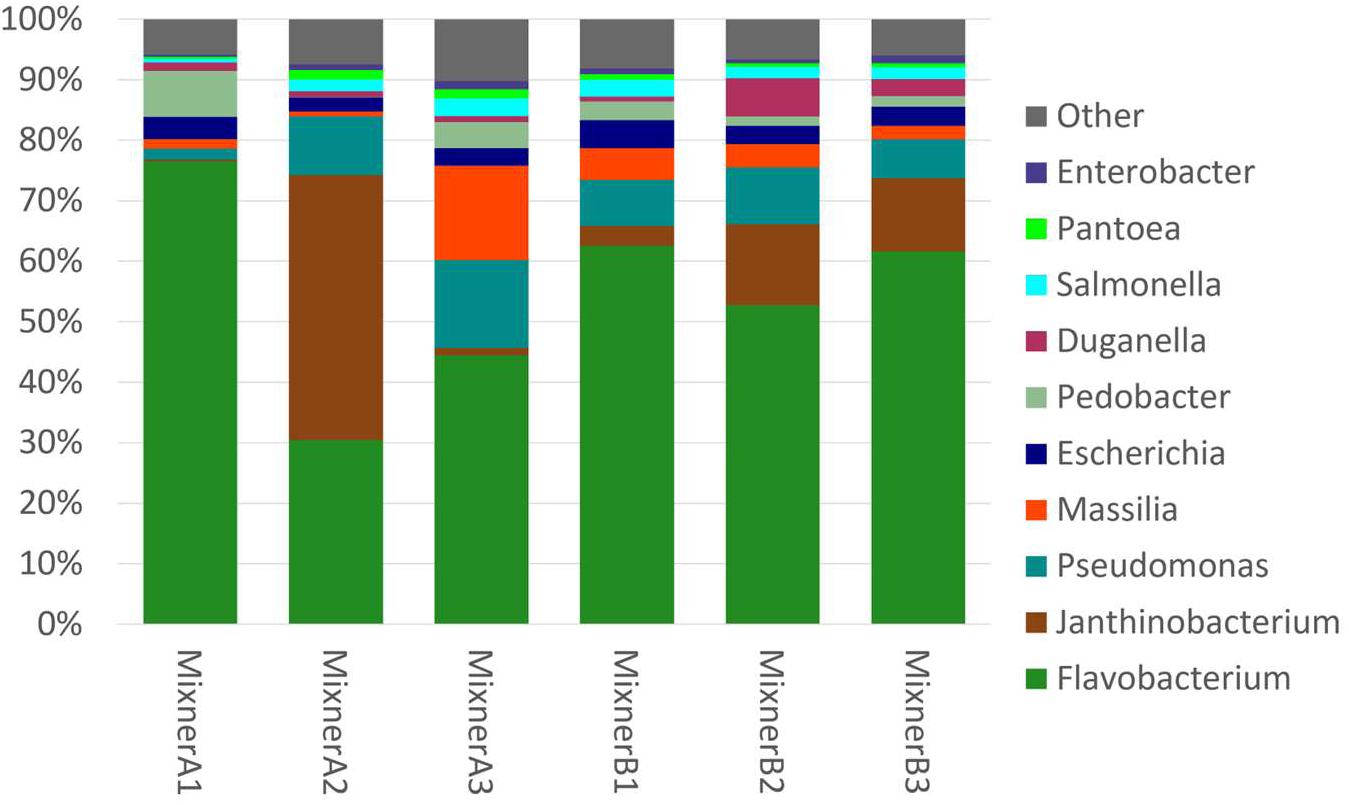
Taxa charts displaying the ten highest abundance bacterial genera from the MixnerA (10% soil suspension, 90% SMC43 overnight, left) and MixnerB (50% soil suspension, 50% SMC43 overnight, right) experimental cultures grown on 0.1X Difco plates for three days at room temperature. Genera are depicted in abundance order. For the MixnerA, an average of 83.3% of the reads were from SMC43 and in MixnerB, an average of 85.8% of the reads were from SMC43. SMC43 reads were removed so that the other bacteria on the plates could be revealed more clearly. Triplicate technical replicates are shown for each treatment.

**Figure 4.**
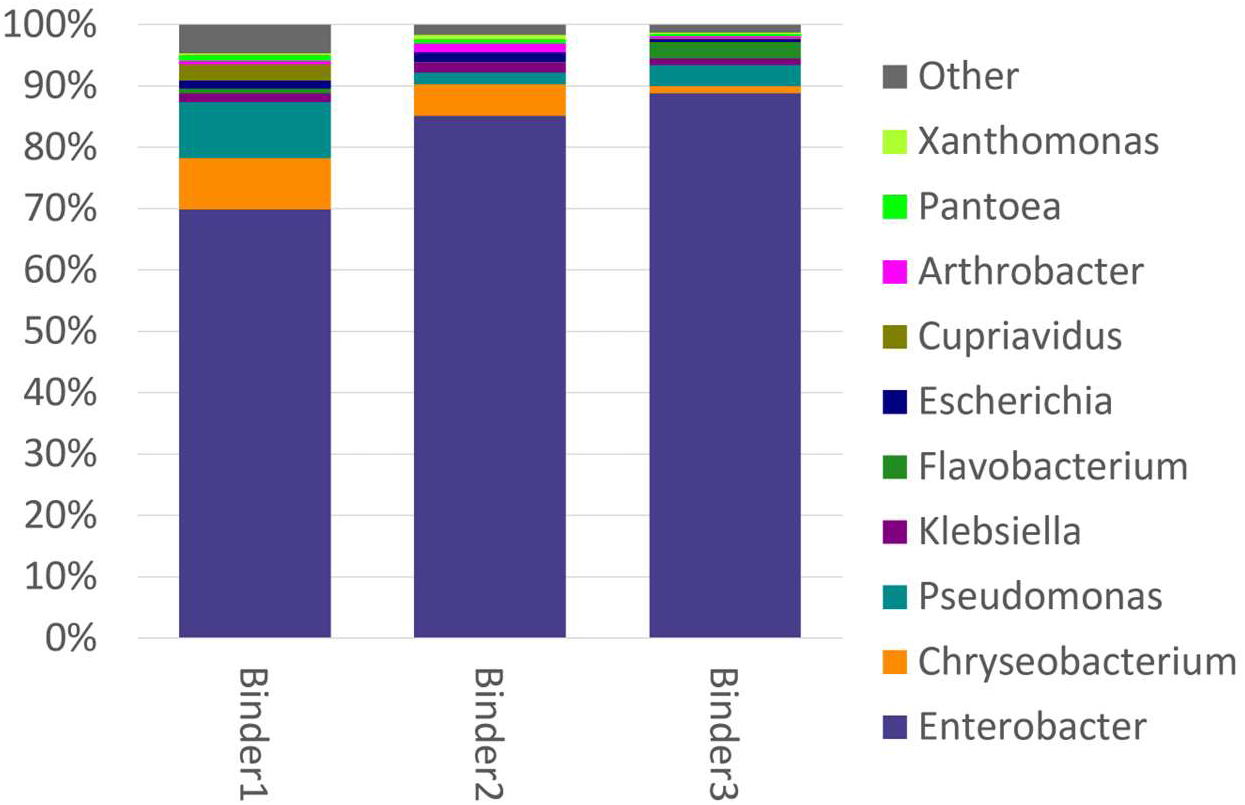
Taxa charts displaying the ten highest abundance bacterial genera that bound to the applicator stick in the absence of SMC43 when grown on 0.1X Difco plates for three days at room temperature. Genera are depicted in abundance order. Triplicate technical replicates are shown for each treatment.

**Figure 5.**
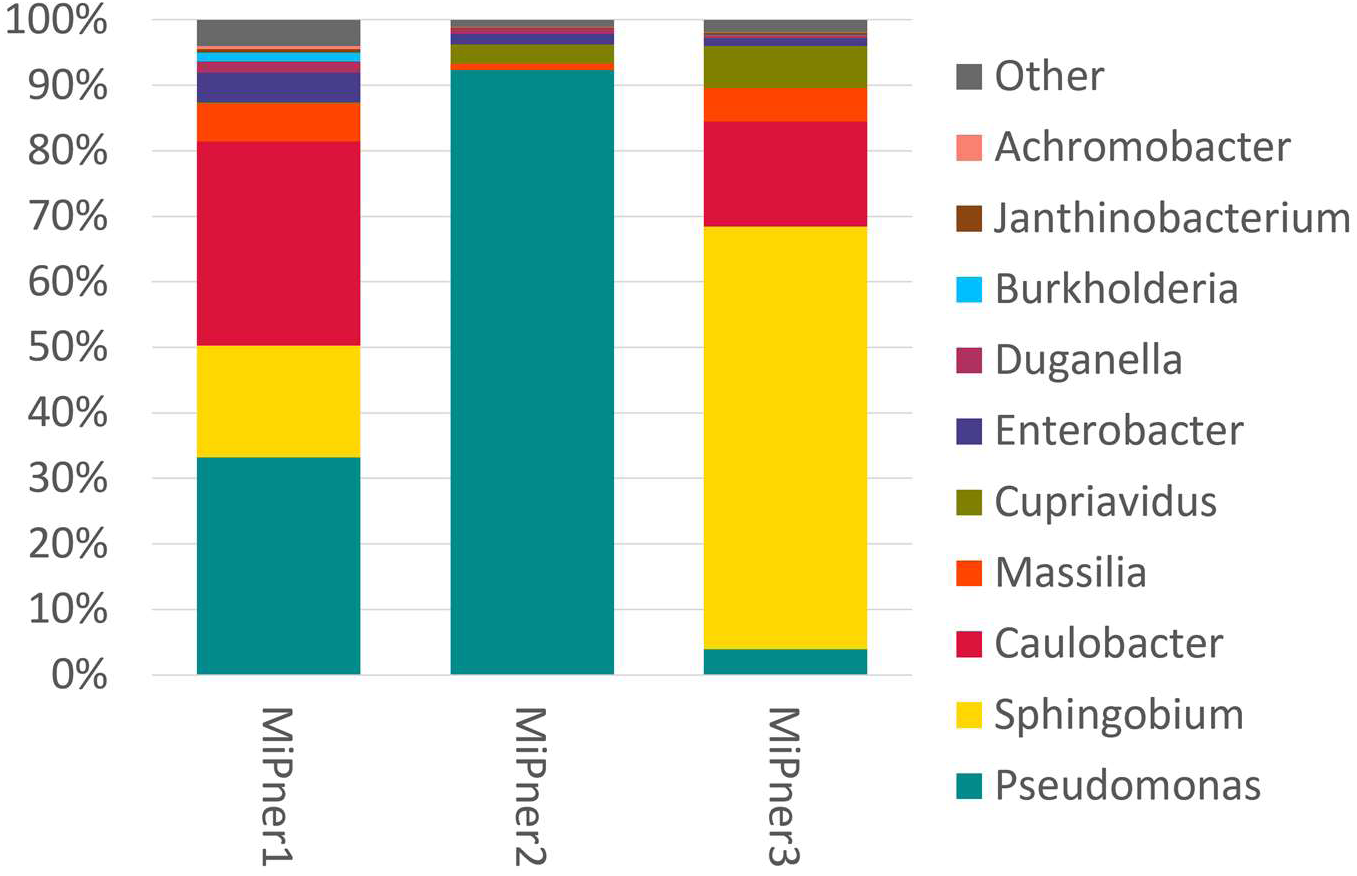
Taxa charts displaying the ten highest abundance bacterial genera observed from the MiPner experiments using the same SMC43 bait and different aliquots of the same soil solution when grown on 0.1X Difco plates for three days at room temperature. SMC43 reads were removed so that the other bacteria on the plates could be revealed more clearly. Triplicate technical replicates are shown for each treatment.

In order to characterize particular MiPners, we picked several of the colonies that were growing on top of the SMC43 streak (e.g., Fig. 1, line 4), for additional streaking on 0.1X Difco plates. The plates from such colony picking indicated a segregation of red and tan colonies. For two of these tan colonies, we sequenced their genomes as described for the sequencing of SMC43. The sequences indicated that one was a strain of *Pseudomonas monteilii* with a 6.3 Mb genome and the other was a strain of *Enterobacter asburiae* with a 3.99 Mb genome. Although *P. monteilii* was not one of the *Pseudomonas* species that bound strongly to the applicator stick, *E. asburiae* was a strong stick binder (Fig. 4), so it seems likely that the *E. asburiae* that we isolated was not actually a true MiPner that bound strongly to SMC43.

### 3.6 Additional MiPner experiments from different soil sources and different times of year

Because soil microbiomes change dramatically over time (28, 29) (and depending on the soil type (30–33) and plant root types/abundances (34–36), we performed two additional MiPner experiments with soil collections from different sites, but using SMC43 as bait. Our hypothesis was that the MiPners that bound to SMC43 would be different in these different soil samples. The same technique was pursued as in the first experiment, and 24 colonies on the SMC43 streak (as in Fig. 1, line 4) were picked and streaked onto 0.1 Difco plates. Of these, 22 picked MiPners yielded red and tan colonies. Two of these tan colonies, from different initial soil collections, were purified and sequenced. One was found to be an ecotype of *Stenotrophomonas maltophilia* with a 4.63 Mb genome and the other was found to be a novel *Mitsuaria* species, with a 3.62 Mb genome, that was most closely related to *Mitsuaria chitinivorans*.

The other two picked MiPners only yielded red solo colonies, but continued to show mixed red/tan bacterial growth in the denser streaking that did not separate out solo colonies. We attempted to sequence one of these, called 3B1D, presumably a mixture of SMC43 and the MiPner, using Oxford Nanopore Technologies platforms. This DNA sequence analysis of 3B1D indicated that there were both one *Mitsuaria* and one SMC43 genomes present, suggesting that this *Mitsuaria* could not grow on this plate type, under our conditions, in the absence of its SMC43 partner. This *Mitsuaria* appears to be a previously uncharacterized species that is most closely related to *Mitsuaria nodu*.

## 4 DISCUSSION

One of the most challenging problems in the study of microbe-microbe interactions in the real world is that we neither understand the micro-environments in which these interactions occur nor have we even a faint idea of the dynamics and depths of involvement of different biological participants. Attempts to create synthetic communities are praiseworthy, but are not proven to actually replicate interactions that occur in nature (10–13). We decided to take a different starting point, an apparent interaction, and create a system that works from that beginning. Obviously, once a two-component community is generated and investigated, adding additional components (established, for instance, by seeing what uniquely binds to a pair of interacting MiPners) will be feasible (37).

### 4.1 The challenges of growing soil bacteria

As has been heavily documented (38–42), most soil bacteria have been (so far) recalcitrant to growth on plates, even though many plate types and growth conditions have been tested. Our experiments growing soil bacteria on 0.1X Difco plates at room temperature under aerobic conditions indicated a great depletion of acidobacteria and actinobacteria, and a great over-representation of proteobacteria, as has been frequently shown by others (38). We expect that different plating conditions would yield different enrichment/depletion patterns (37, 43, 44). Why these microbes are recalcitrant to culturing is not known, but it is expected to be associated with an unknown necessary component or components of their micro-environments (43). Perhaps one of these unknown components is a microbial partner (45), as manifested in our SMC43-*Mitsuaria*-3B1D result. Syntrophic interactions are not uncommon in microbial networks, of which this interdependence and need for co-culture may be an example (47). We have seen other such “cannot grow without the bait” examples from MiPner experiments with other bait species (unpublished), so it is possible that MiPner technology may be one general tool for future isolation of such recalcitrant microbes.

Of course, our SMC43-associated *Mitsuaria* may be able to grow on some plate types if we pursued a full round of investigations. Many *Mitsuaria* grow on several different plate types, as we have seen in our lab, including 0.1X Difco plates (46), but we believe it is likely that many such future “bait-requiring” microbe isolations will be of species that have an absolute partner requirement. Regardless, the SMC43-*Mitsuaria*-3B1D interaction on 0.1X Difco plates indicates a pairwise interaction that is obligate for this *Mitsuaria*’s growth, and future studies showing what SMC43 provides in this relationship will be of great interest.

### 4.2 MiPner specificity

The different and taxonomically limited set of MiPner microbes identified, compared to our starting soil and to other selective steps (e.g., plate type) in the technique, indicates that the microbe-microbe binding is highly selective and robust enough to avoid removal by a simple rinsing step. Moreover, this binding requires only a few minutes to generate this specificity and durability. Preliminary studies in our lab using different bait species on the same soil suspension (unpublished results) have suggested that each bait generates a separate set of MiPners that are enormously enriched at the genus level. Once pursued, we expect that observed species-level binding specificities and enrichments will be even more dramatic. Hence, there is an unlimited potential for using MiPner technology to find potential interacting partners with most other microbes, and this should extend beyond just bacteria.

Isolation of colonies growing on the bait streak is not likely to only yield MiPners, as shown with our *E. asburiae* result. Anything that binds the applicator stick found in the study does not need to bind the bait, although it is required to grow with the bait on the plate type used. For this study, we would be confident of the MiPner status of any *Sphingobium* or *Caulobacter* isolated, which could be further confirmed by reciprocal binding studies.

### 4.3 The full set of potential bait partners

There is no reason to believe that any bacterium interacts with the same set of microbial partners in all environments. Our investigations of different soils with SMC43 indicated different final MiPner outcomes, dependent on soil source. The two *Mitsuaria* that we found in a subsequent experiment to the one described in detail here, represent a genus that was fully absent from all our sequencing in the first experiment. Hence, discovery of the full set of SMC43 MiPners that may be real-life microbial partners would be best pursued with a number of different soil sources. And this will be equally true for any other microbe used as bait.

### 4.4 Mixner results

We call the community DNA sequence results of growing one bait microbe with a soil suspension of microbes to be the outcome of a Mixed preMiPner, or Mixner, experiment. Our Mixner outcomes indicate variable survival patterns of the soil-solution microbes depending upon the dose of the bait microbe. Of course, SMC43 may provide a severe example of this phenomenon because of its production of prodigiosin, a potent anti-microbial (26). However, many bacteria produce antimicrobials, so we predict this ratio-dependence result to be generally true with any bait microbe or any mixed microbe suspension. Perhaps at lower bait dosages, other microbes have a greater opportunity to build communities that will resist any negative (or positive) contributions from the bait microbe.

### 4.5 MiPner enrichment

Several steps of enrichment led to the isolated microbes that were dubbed MiPners. The choice of a liquid suspension, rather than total soil, as the initial soil microbe source was one such enrichment/depletion step. As noted, growth on a plate and survival of exposure to SMC43 were other selection steps, each with unique outcomes. The primary goal of this technology, and thus the most interesting enrichment for us, is to find paired candidates for a specific SMC43-MiPner interaction. Starting with these functional components of a durable and highly-specific binding, study of these two-species interactions can proceed in a wealth of directions. Characterization of each of these pairs of interactions are warranted by such techniques as annotation for gene-enrichment by the selection process, optical studies of the physical interaction, forward genetic searches for genes that decrease or increase the partnership, reverse genetics of genes likely to be involved in the binding and other interactions, transcriptomic/proteomic/metabolomic analysis of inductions/repressions by the interactions, and many others too numerous to list. All of these are beyond the scope of our current investigation.

It should be noted that none of the controls that we pursued in this study would be necessary to pull out MiPners that bound to the bait. Just the simple bait binding to an applicator, followed by the second immersion and plating, would be sufficient to find a candidate partner. However, the various controls were of value in determining the likelihood that the identified microbes truly were MiPners, especially in distinguishing microbes that bound to the applicator stick without bait involvement. Full confirmation of a MiPner would be best pursued by subsequent studies, especially by using the MiPner as bait to see if it reciprocally pulls out its partner that was the initial bait.

### 4.6 MiPner generality

As should be clear, there is absolutely nothing about the MiPner strategy that is limited to any specific bait microbe or to any specific microbial community. The animal gut (47–49) (Dillon and Dillon, 2004; Gill et al., 2006) or bodies of water (4, 50) and such fascinating microbial worlds as permafrost (51), the digestive fluids of pitcher plants (52, 53) or waste tailings (54) will be equally accessible to this technology. There is potential for viruses, fungi, protists, tiny invertebrates or archaea to be used as baits or followed as MiPners. Moreover, it is difficult to overestimate the speed, simplicity, robustness and low cost of this approach. We hope that many laboratories will join us in MiPner experimentation, so that we can begin to assemble a microbial interaction atlas, starting with two species at a time.

## ACKNOWLEDGEMENTS

The authors would like to thank Molly Levine for her contribution to SMC43 annotation as part of the GENE4240L course at the University of Georgia, USA.

## SUPPLEMENTARY MATERIALS

**Figure S1:**
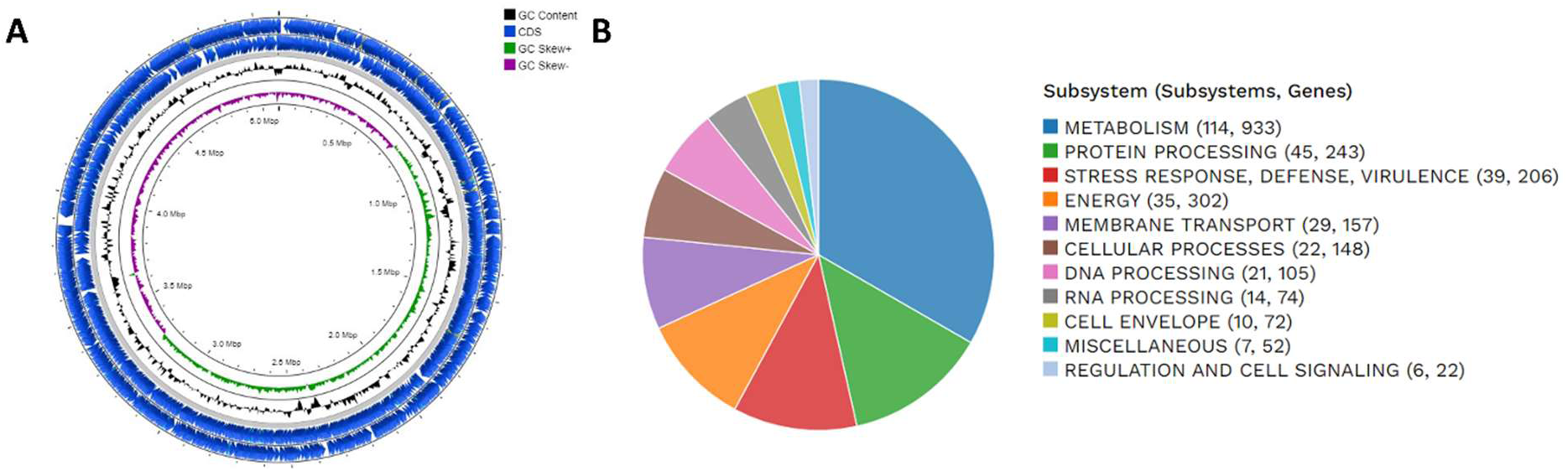
A) Genomic map and BRIG analysis of SMC43 featuring the circular map of the genome. Outer circle to inner circle (CDS, GC content, GC skew+, GC skew -, and GC), B) Subsystem distribution based on RAST SEED analysis of SMC43. The pie chart organizes cellular processes. The number of protein-coding genes predicted to be involved in each cellular process are indicated in parentheses.

**Figure S2.**
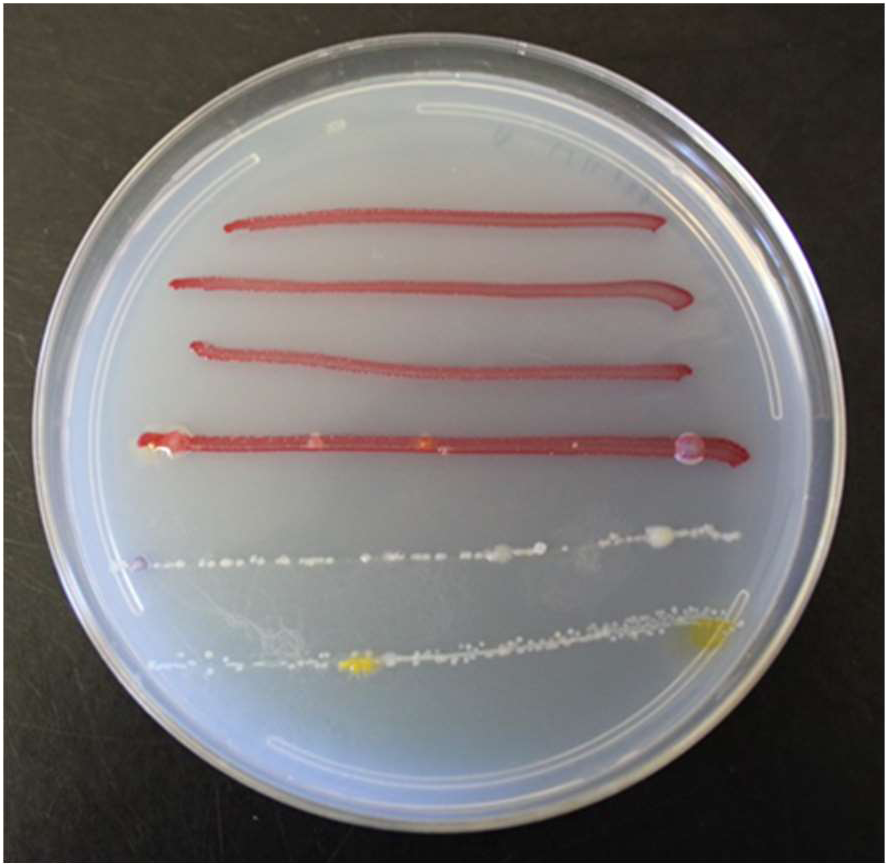
A demonstrative 0.1X Difco plate grown for three days at room temperature with streaks of SMC43 (lines 1, 2), streaks of SMC43 plus putative MiPners (lines 3, 4) and with streaks of microbes that bound to the applicator stick in the absence of SMC43 pre-binding (lines 5, 6).

